# Pupillometry as a reliable metric of auditory detection and discrimination across diverse stimulus paradigms in animal models

**DOI:** 10.1101/2020.11.16.385286

**Authors:** Pilar Montes-Lourido, Manaswini Kar, Isha Kumbam, Srivatsun Sadagopan

## Abstract

Estimates of detection and discrimination thresholds are often used to explore broad perceptual similarities between human subjects and animal models. Pupillometry shows great promise as a non-invasive, easily-deployable method of comparing human and animal thresholds. Using pupillometry, previous studies in animal models have obtained threshold estimates to simple stimuli such as pure tones, but have not explored whether similar pupil responses can be evoked by complex stimuli, what other stimulus contingencies might affect stimulus-evoked pupil responses, and if pupil responses can be modulated by experience or short-term training. In this study, we used an auditory oddball paradigm to estimate detection and discrimination thresholds across a wide range of stimuli in guinea pigs. We demonstrate that pupillometry yields reliable detection and discrimination thresholds across a range of simple (tones) and complex (conspecific vocalizations) stimuli; that pupil responses can be robustly evoked using different stimulus contingencies (low-level acoustic changes, or higher level categorical changes); and that pupil responses are modulated by short-term training. These results lay the foundation for using pupillometry as a high-throughput method of estimating thresholds in large experimental cohorts, and unveil the full potential of using pupillometry to explore broad similarities between humans and animal models.

## Introduction

Even in the absence of luminance changes, pupil size fluctuates in response to a variety of endogenous and exogenous factors. In relaxed human subjects, the pupil diameter (PD) exhibits sustained low-frequency oscillations [1] mainly resulting from modulation of parasympathetic neural activity [2,3]. With task engagement, the pupil dilates with increasing arousal or increasing mental effort [4–7]. Increased effort from the subject is necessary, for example, when searching for visual targets [8,9], or listening for meaningful sounds in cluttered, ambiguous, or noisy acoustic conditions [10–13]. These PD changes can be attributed to endogenous mechanisms, i.e., when the subject regulates their arousal or attention to achieve better task performance [5,10]. In animal models, PD is similarly modulated by listening effort and other top-down factors, and PD strongly correlates with cortical activity states [14–18], and has been shown to closely track noradrenergic and cholinergic tone at fast and slow timescales respectively [19]. Pupillometry can thus be used as a window into spontaneously fluctuating neuromodulatory tone across waking and sleep states [20,21], providing a non-invasive metric to evaluate animal models of neuromodulatory defects in clinical populations [22].

PD changes can also be evoked exogenously by providing stimuli that capture the subject’s attention in a bottom-up fashion. This could be accomplished, for example, by providing an occasional unexpected stimulus [7,18,23,24], a salient or familiar stimulus [25], or by abruptly changing the statistical distribution of stimuli [23,26–29]. One such example is the auditory ‘oddball’ paradigm, where the subject is habituated to a standard, predictable stimulus pattern, and is ‘surprised’ by infrequent deviant stimuli [23,24,26,29]. We emphasize that PD changes triggered by the deviants reflect a change in the internal state of the subject, and are unrelated to visual processing. Because PD data can be obtained non-invasively from untrained human and animal subjects using the oddball paradigm, it is well-suited for comparing auditory detection and discrimination between humans and animal models. Combining pupillometry with other non-invasive measurements such as EEG recordings, broad similarities between human and animal subjects could be established, which can then be followed up in animals using invasive recordings to interrogate underlying neural mechanisms. Because changes to pupil responses, possibly corresponding to perceptual difficulties, occur in some clinical populations [30–32], these signals also hold promise as phenotypes for potential animal models of these disorders. However, past pupillometric estimates of auditory detection and discrimination thresholds in animals have been restricted to simple stimuli such as pure tones. Previous studies have also not examined what other stimulus contingencies affect PD changes, and if PD changes can be shaped by experience or learning (but see ref. [33]).

In this study, we explored various factors affecting sound-evoked PD changes in detail. We demonstrate that pupillometry can be used to estimate detection and discrimination thresholds across a range of simple (tones) and complex (conspecific vocalizations) stimuli; that PD changes can be robustly evoked using different stimulus contingencies (acoustic changes or categorical changes); and that PD changes are modulated by short-term training. These results lay the foundation for using pupillometry as a high-throughput method of estimating auditory detection and discrimination thresholds in large experimental cohorts, and unveil the full potential of using pupillometry to explore auditory similarities between human and animal models.

## Results

We obtained PD data from eleven head-fixed guinea pigs (GPs; three to seven animals per paradigm with overlap) over a range of simple (detection of changes in fundamental frequency (F0)) and complex (detection of an auditory figure embedded in a random background) auditory ‘tasks’. Tasks were structured as auditory oddball paradigms, where a standard stimulus was presented >90% of the time, interspersed randomly with deviant stimuli whose acoustic distances from the standard stimuli were parametrically varied. To minimize the effects of animal adaptation over the duration of a session, we restricted each pupillometry session to contain few deviant trials, and restricted the total time spent in pupillometry experiments to ∼30 minutes per day. Deviants occurred in a pseudorandom order determined by a Latin square design (see Methods). In normal illumination conditions, GP pupils are quite dilated (∼4.5 mm dia.), leaving little dynamic range for us to observe further sound-evoked dilations. Therefore, to ensure sufficient dynamic range as well as consistent baselines across sessions and subjects, we illuminated the eyes with diffuse white light to bring baseline PD to about 3.5 mm, and kept illumination levels constant over the experimental session. During each session, we monitored animal motion (postural shifts) via a piezoelectric sensor and imaged the pupil using infrared illumination (Fig. 1a). Under these experimental conditions, we could reliably elicit pupil dilation responses to deviants using a wide variety of stimuli.

**Figure 1:**
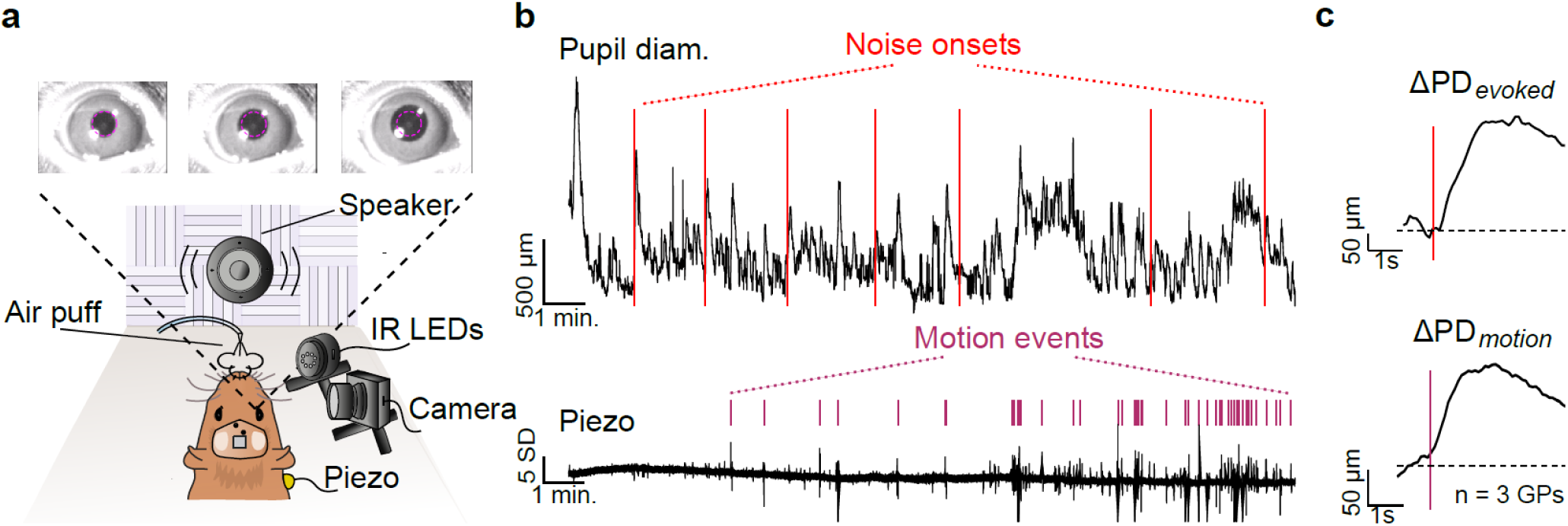
Stimulus-evoked and motion-related PD changes. **(a)** Pupillometry setup showing video frame captures of sound-evoked pupil dilation. Magenta circle corresponds to baseline PD. **(b)** Representative pupil diameter trace (top) and output of piezoelectric sensor (bottom) from a single experimental session. Red lines correspond to stimulus times. Purple ticks correspond to automatically detected motion events. **(c)** Stimulus-triggered (top) and motion-related (bottom) changes in PD (ΔPD) averaged across 3 animals. Red line corresponds to stimulus onset; purple line corresponds to detection of motion event. Note that the onset of pupil dilation precedes motion onset.

### Time-course of stimulus-evoked and motion-related pupil responses

We first determined the time course of stimulus-evoked pupil responses by presenting 1 s-long noise bursts occurring once every ∼1.5 minutes. Figure 1b (top) illustrates an example PD trace acquired over the duration of an entire session (∼12 minutes). Figure 1b (bottom) represents the output of a piezoelectric sensor placed beneath the animal. We observed that the pupil trace was in a state of constant flux, changing 1) in the absence of any exogenous, experimenter-provided stimuli, 2) in response to the noise bursts (red lines), and 3) around periods of animal motion, detected using an automated algorithm (see Methods). When we averaged the pupil trace triggered on stimulus onsets (Fig. 1c, top), we obtained a response with ∼300 ms latency, reaching a peak dilation of ∼7% increase from baseline occurring ∼1.8 s after stimulus onset, and slowly returning to baseline (after ∼5 s of stimulus offset). Consistent with previous studies, when we averaged the pupil trace triggered by motion events (Fig. 1c, bottom; averaged over 3 subjects), we obtained a similar pupil time course, except that dilation onset preceded the detected motion event by about 0.5 s [15,17,18]. Thus, to avoid confounding stimulus-evoked and motion-related pupil dilation, stimuli that were presented within 7 s of a motion event were discarded from further analyses (see Methods). Note that while our piezo sensor-based motion detection picked up postural shifts and limb movements, we did not detect orofacial movements (whisking, licking, or chewing) that are also associated with PD changes [34]. Thus, it is possible that some stimulus-driven pupil responses also contain some of these motion-related PD changes. Automated motion detection from videography could provide higher accuracy in excluding motion effects from pupil traces.

In the context of an auditory oddball paradigm, we separately averaged standard and deviant trials. For example, in Figs. 2b and 2c, we plot average PD changes to standard stimuli (white noise bursts) and deviant stimuli, which were GP vocalizations (purr calls; Supplementary Fig. 1) recorded in our animal colony. The GP pupil dilation time course was strikingly similar to previously published pupil time courses of humans using an auditory oddball paradigm (Fig. 2a; reproduced with permission from [26]). Across 3 subjects, the average time courses of pupil dilation triggered by auditory deviant stimuli (Fig. 2c), motion events (Fig. 2d), and by the delivery of a 100 ms air puff (Fig. 2e) were similar. These data suggest that the overall time course of pupil dilation reflects non-specific underlying mechanisms. Stated differently, the pupil response could be conceptualized as a pupil impulse response function that could be convolved with known motion event times and stimulus times to result in a pupil trace estimate, with only the weighting being different between stimulus types (note that the residual could contain pupil dilation events arising from unknown factors, analyzed later in the manuscript). However, this was true only for stimuli with positive weights, or sound onsets. When we measured the average pupil response triggered by 1 s-long silent gaps embedded every ∼1.5 minutes in a constant noise background, we observed a pupil constriction rather than a dilation (Fig. 2f). In fact, pupil dilation occurred at the offset of the gap when the continuous noise resumed. Thus, pupil dilation with a stereotypical impulse response seems to occur in response to sound presence or positive changes, but not in response to silence or negative changes.

**Figure 2:**
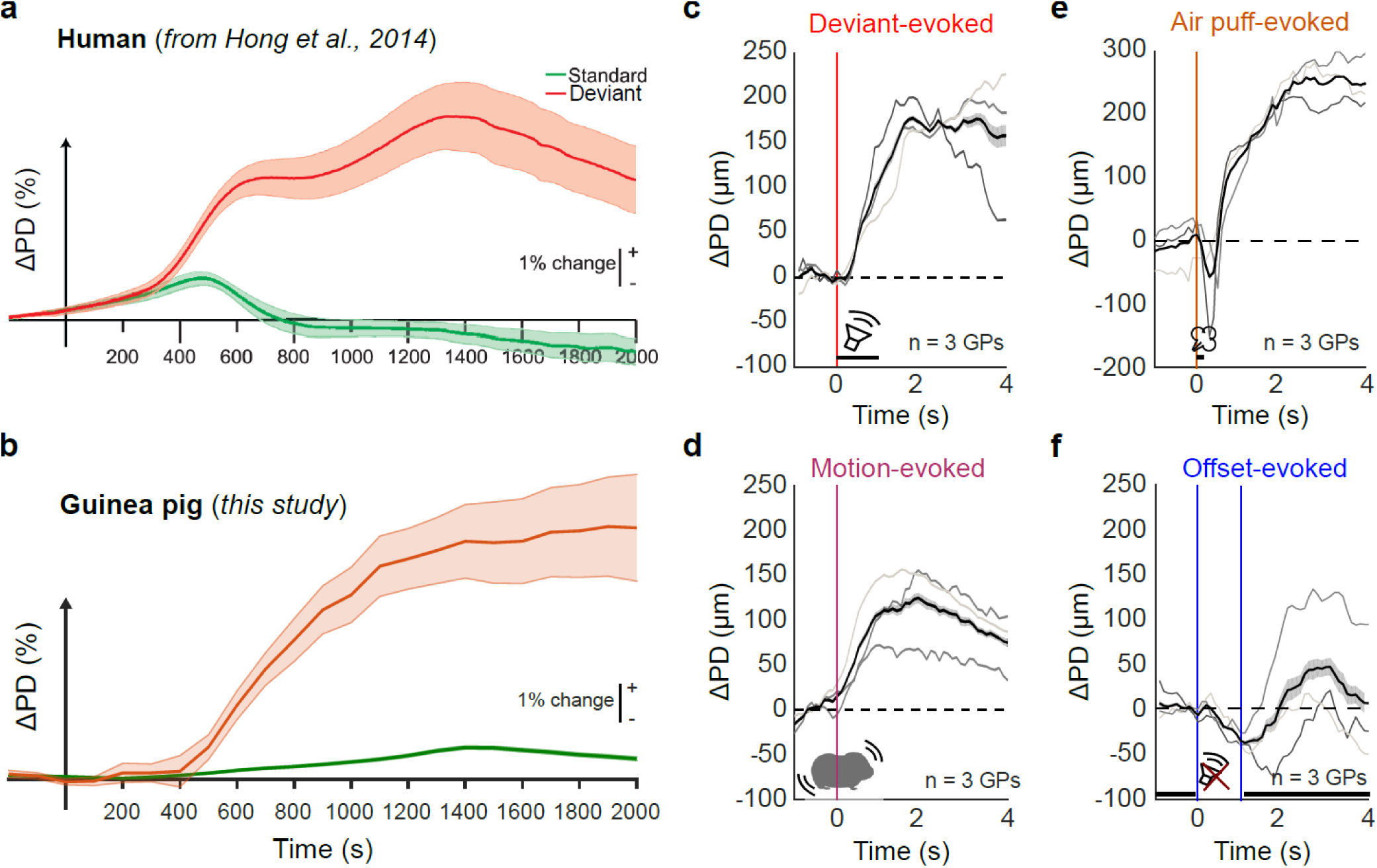
Time course of PD changes evoked by different stimulus conditions. **(a)** PD changes evoked by standard (green) and deviant (red) stimuli in an auditory oddball task in humans, adapted with permission from Hong et al., 2014. **(b)** PD changes evoked by standard (white noise) and deviant (GP call) stimuli in GPs, showing a similar time course to that observed in humans. Shading corresponds to ±1 s.e.m. **(c – e)** Similar average time courses of PD changes were evoked by auditory deviant stimuli **(c)**, animal motion **(d)**, and a brief (100 ms) air puff **(e). (f)** In contrast, no pupil dilation was evoked when a 1 s long gap was embedded in continuous white noise. Rather, pupil dilation was evoked when continuous noise restarted following this gap. Vertical colored lines correspond to stimulus onsets, gray lines correspond to data from individual subjects, black lines correspond to means across subjects, shading corresponds to ±1 s.e.m.

### PD changes are proportional to the acoustic distance between the standard and deviant

We next asked whether the magnitude of the pupil response scales with the acoustic distance between deviant and standard stimuli. As a simple case, we tested whether we could quantify the ability of GPs to discriminate harmonic complexes of different fundamental frequencies (F0; Fig. 3a). We presented an oddball sequence where standards were harmonic complexes with F0 = 440 Hz, and deviants that were harmonic complexes with F0s that were higher than the standard F0 by a ΔF ranging from 0.25 to 2.0 semitones (a semitone equals 1/12^th^ of an octave; Fig. 3b). While the standard evoked little pupil dilation, we observed robust pupil responses to most of the deviant stimuli (single subject example in Fig. 3c, average over 7 GPs in Fig. 3d). The average change in pupil diameter (ΔPD), measured in a window ranging from 1 to 2 s after stimulus onset, appeared to scale linearly with ΔF (Fig. 3e; but note that ΔF is in logarithmic units of semitones), as did the fraction of trials whose peak pupil response differed significantly from the distribution of standard peak pupil responses (Fig. 3f). The ΔPD was statistically significant for ΔF values of 0.75 semitones and above (Supplementary Table 1).

**Figure 3:**
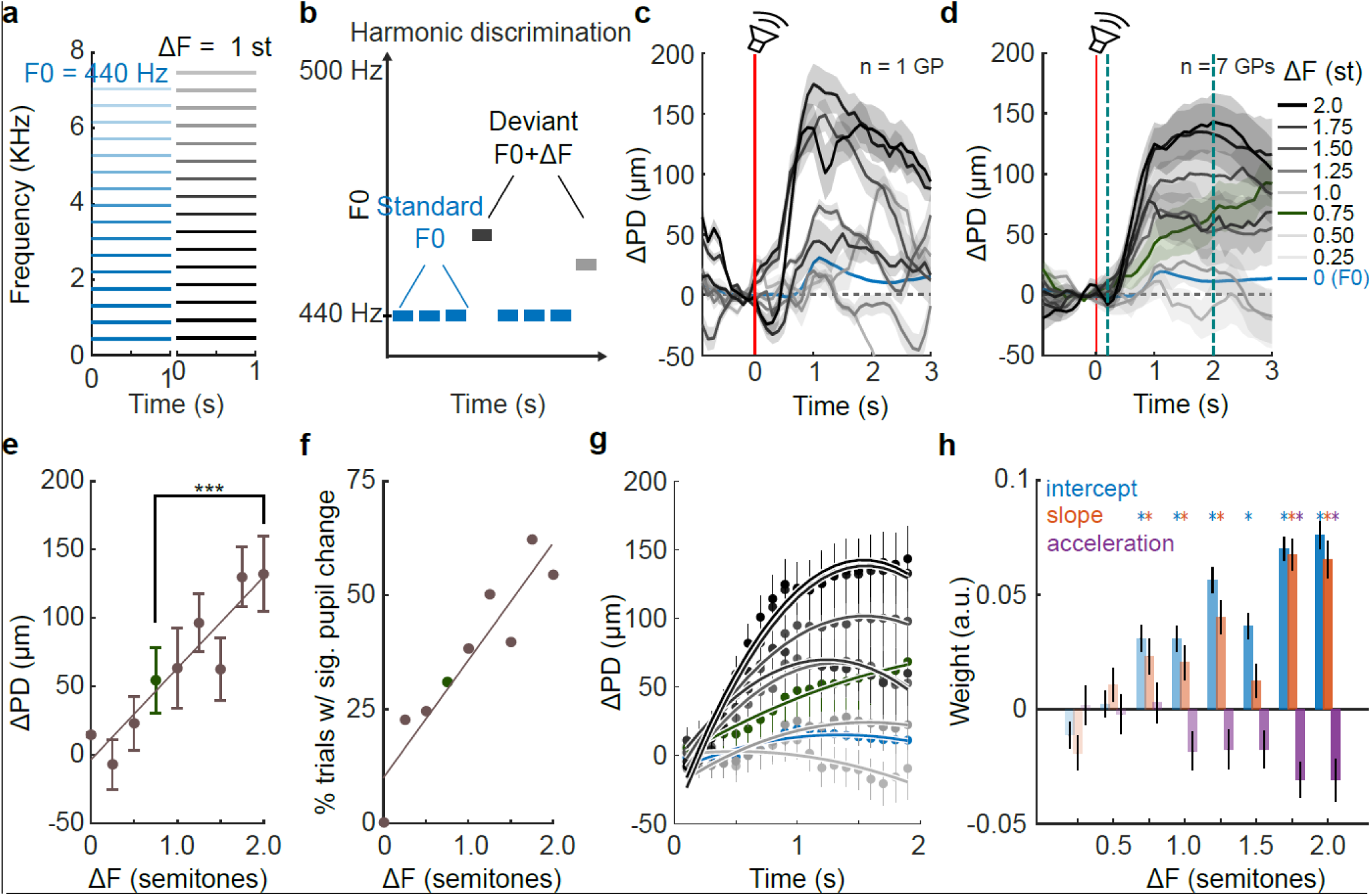
Pupillometric estimate of harmonic tone discrimination thresholds. **(a)** Schematic spectrograms of standard (blue) and deviant (gray) stimuli. Color intensity corresponds to amplitude of each frequency component. **(b)** Schematic of the oddball paradigm used to probe harmonic tone discrimination. **(c)** PD changes evoked by deviant stimuli in one subject. Color gradient corresponds to the F0 difference between the deviant and standard F0s. Lines correspond to mean PD and shading corresponds to ±1 s.e.m. Red line corresponds to stimulus onset. **(d)** Same as **(c)** but pooled and averaged over 7 subjects. Green line and shading corresponds to pupil trace at threshold (ΔF = 0.75 semitones). Teal dashed lines correspond to the GCA window. **(e)** Average PD changes computed in a 1-s window beginning 1 s after stimulus onset as a function of deviant F0 (whiskers correspond to ±1 s.e.m.). Significant changes in pupil diameter were observed for deviants with F0s 0.75 semitones or greater than the standard F0 (green; ***: p<0.0001, ANOVA followed by Bonferroni-corrected post-hoc tests; exact p-values in Supplementary Table 1). Line is linear fit. **(f)** The percent of trials (circles) in each deviant condition that showed a significant pupil dilation compared to standard stimuli. Line is linear fit. **(g)** GCA fit to the rising phase of PD changes (teal dashed lines in **(d)**). Dots are mean pupil diameter in 100 ms time bins, whiskers correspond to ±1 s.e.m. Solid lines correspond to mixed-effects model fits. Colors as in **(d). (h)** GCA weight estimates. Colors correspond to the weights of the intercept (blue), slope (red), and acceleration (purple) terms used to fit the PD traces. Asterisks denote statistically significant regression weights (exact p-values in Supplementary Table 2). Significant differences in pupil traces were observed for deviants with F0s 0.75 semitones or greater compared to standard F0s.

The scaling of the pupil response with acoustic distance could result from two possible trial-wise response distributions: 1) trial-wise responses could be normally distributed with gradually scaled means for each deviant, and the maximum responses for each deviant could also scale with distance, or 2) trial-wise responses could be all-or-none with a similar magnitude of change for all deviants, but with the fraction of trials eliciting a response changing for each deviant. In this case, the trial-wise distributions would be bimodal, and maximum responses for each deviant would be similar. Analysis of the pupil traces supported the former model – maximal responses for each deviant were also scaled, and trial-wise responses were not bimodal – indicating that the pupil response magnitude indeed scaled with acoustic distance.

To further quantify and statistically evaluate PD changes, we used growth curve analysis (GCA; see refs. [13,35]) to fit a mixed-effects model consisting of subject-level intercepts as random effects, and up to quadratic-time orthogonal polynomials as fixed effects, to the rising phase of the pupil response (Fig. 3g, see Methods). The weight estimates of the intercept, linear, and quadratic fixed effects terms are plotted as a function of stimulus distance in Fig. 3h. Hypothesis tests for linear regression model coefficients revealed that compared to the standard stimulus, pupil responses of deviants with a ΔF of 0.75 semitones or greater were significantly different (Supplementary Table 2). From these analyses, we concluded that the pupillometric estimate for F0 discriminability in GPs at F0 = 440 Hz was about 0.75 semitones (= 20 Hz, or 4.5% change).

As mentioned earlier, the pupil diameter is in a state of constant flux, influenced by unobserved endogenous factors as well as exogenous factors. To determine the extent to which stimulus-related changes explained the overall occurrence of pupil dilation events, we first detected all pupil dilation events occurring over the course of an experimental session (see Methods). We then determined the fraction of these events that occurred within 2 s of stimulus presentation or a motion event, and could thus be associated with observed endogenous (motion) or exogenous (stimulus-driven) factors. Over the seven GPs used in this experiment, we observed on average 161 ± 15 pupil dilation events during an experimental session, of which 79% ± 4% were associated with a stimulus or motion event.

Next, we asked if pupillometry could be used to estimate thresholds for detecting conspecific vocalizations (calls) in white noise. Framed as an oddball paradigm, we used white noise bursts as standards and noisy calls at different signal-to-noise ratios (SNRs) as deviants (Figs. 4a – c). From pilot experiments, we chose SNR values to maximize sampling in the linear part of the psychometric curve. Deviants were randomly selected from a set of 8 exemplar calls (r.m.s.-normalized) belonging to the same category, and white noise added to reach the desired SNR value. We determined detection thresholds for two different GP call types, *chuts* and *purrs* (Supplementary Fig. 1). As a control for behavioral relevance, we also obtained detection thresholds for harmonic complexes in noise (constant F0 = 440 Hz). In these experiments, the standard stimuli (white noise bursts) only evoked small PD changes that did not significantly differ from the baseline. In contrast, deviant stimuli (calls in noise) evoked large and robust PD changes. In Supplementary Video 1, we show an example snippet of the pupil response to a high-SNR call deviant. Both the response magnitude (Fig. 4d – single subject, and Fig. 4e – average of 5 subjects) and the fraction of statistically significant trials (Fig. 4g) scaled with SNR. The fraction of significant trials at each SNR was well-fit by a psychometric function (see Methods).

**Figure 4:**
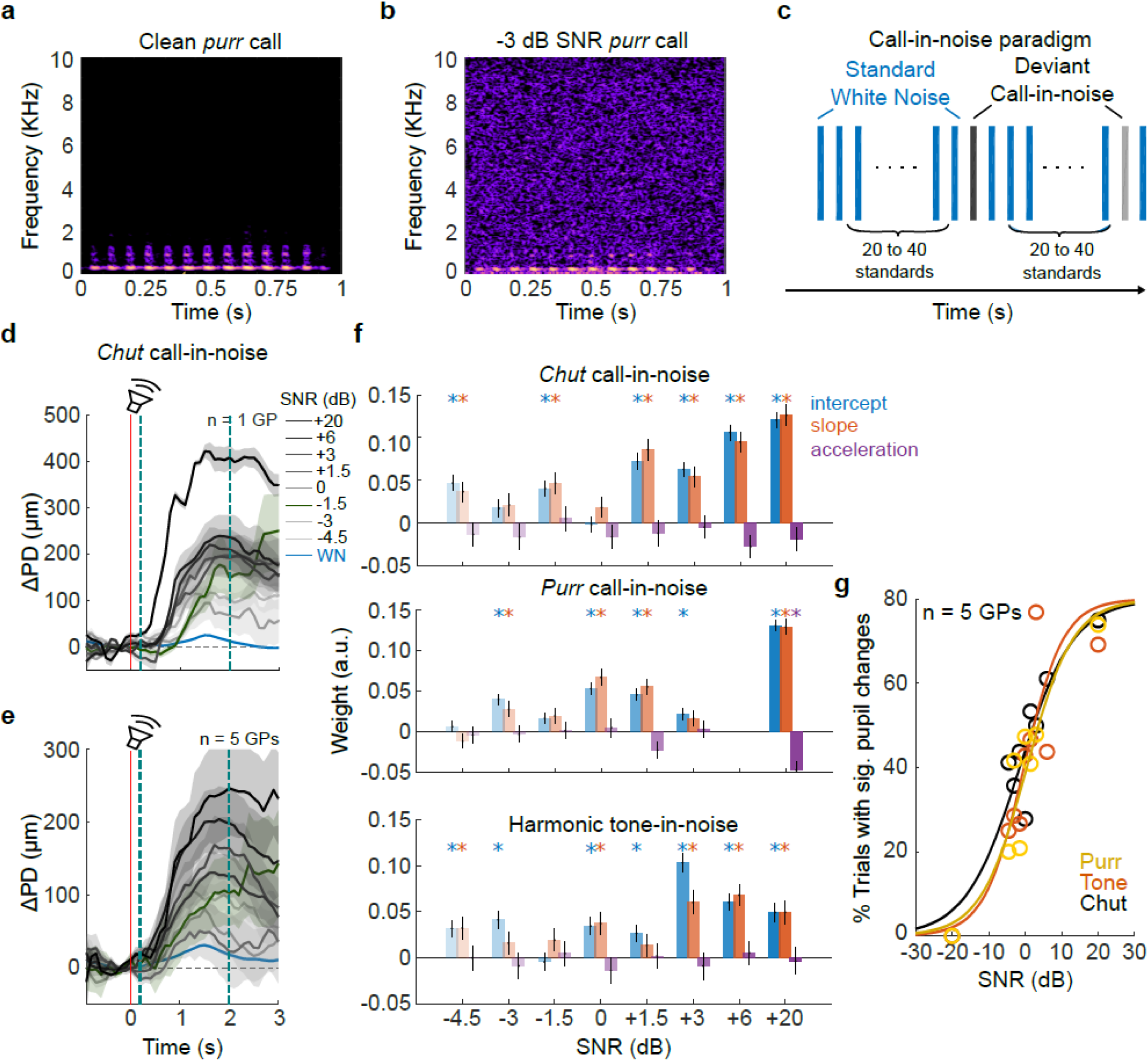
Pupillometric estimates of call-in-noise detection thresholds. **(a)** Spectrogram of a guinea pig *purr* call. **(b)** *Purr* call at -3 dB SNR, obtained by adding white noise to the ‘clean’ *purr* call in (A). **(c)** Structure of call-in-noise detection paradigm. Standard stimuli were white noise bursts; deviants were calls embedded in white noise at different SNR levels. **(d)** Examples of PD traces from one subject for detecting *chut* calls in noise. Blue line and shading corresponds to pupil responses to standard stimuli, and black line and shading correspond to mean ±1 s.e.m. of pupil responses to deviants. Color gradient corresponds to stimulus SNR. Green line and shading denote pupil trace at threshold (−1.5 dB SNR). Red line signifies stimulus onset, and teal dashed lines demarcate the GCA window. **(e)** Pooled and averaged pupil responses across 5 subjects **(f)** GCA weight estimates for the detection of *chuts, purrs*, and harmonic tones in noise (bar colors as earlier). Weight estimates for deviants were significantly different from the standard for most SNRs above -1.5 dB SNR. Asterisks correspond to p < 0.01 (hypothesis test for linear model coefficients, exact p-values in Supplementary Tables 3 – 5). Note that the +6 dB SNR data point was not sampled for *purr* calls. **(g)** Psychometric functions fit to the percent of trials that evoked significant PD changes as a function of SNR for *purr*-in-noise (yellow), *chut*-in-noise (black), and harmonic tone-in-noise (orange). Psychometric functions were largely similar, reaching 50% of maximum at about -1.5 dB SNR.

Using GCA, we determined that most pupil changes evoked by deviants of -1.5 dB SNR or greater were statistically significant (Fig. 4f; Supplementary Tables 3 – 5). Results were similar across both call categories as well the harmonic complex (Fig. 4g). Defining the call-in-noise detection threshold as the SNR necessary to reach the half-maximum of the psychometric curve, we determined that for all stimulus types, the detection threshold was about -1.5 dB SNR.

### Pupil responses are modulated by air puff training

Three factors limited the number of trials acquired per experimental session: 1) the animal adapted to the deviants and responses progressively weakened, 2) overall arousal trended downward over the session, and 3) for some animals, motion increased gradually resulting in an increasing number of discarded trials. To mitigate these issues and to maintain sustained arousal levels for longer time periods, we modified the auditory oddball paradigm to additionally deliver an air puff to the animal’s snout after each deviant stimulus (Fig. 5a). The delay between deviant onset and the air puff was set at 2.5 s to allow the deviant-triggered pupil response to reach a peak before the onset of the air puff. To test if we could indeed sustain high arousal levels for longer time periods, we increased the number of deviants per session from 8 to ∼48. Sessions were repeated for 10 days, which we divided into three phases for analysis: early (Days 1 – 4), middle (Days 5 – 7), and late (Days 8 – 10). Average pupil responses (from 3 GPs) to the standard (white noise) and deviant stimuli (*purr* call in noise at different SNRs) during each of these phases is shown in Fig. 5b.

**Figure 5:**
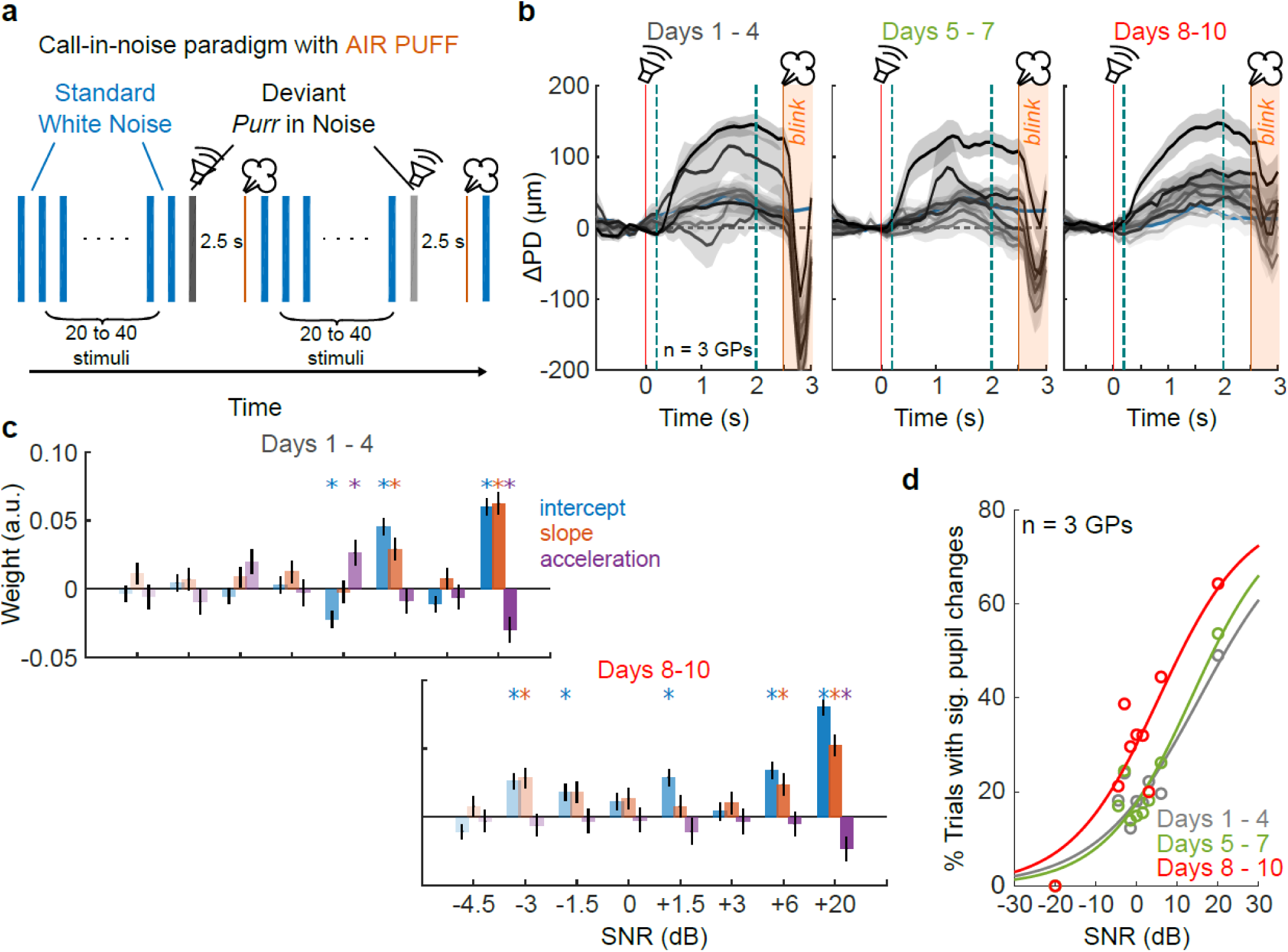
Air puff training increases reliability when using pupillometry to probe call-in-noise detection. **(a)** The paradigm used was similar to Figure 4, except that brief air puffs (orange lines) were delivered 2.5 s after each deviant onset. **(b)** Pupil responses of 3 animals as they progressed through 10 days of air puff training. Color gradient corresponds to deviant SNR, lines and shading correspond to mean ±1 s.e.m. of deviant pupil responses. Orange line is air puff onset; orange shading corresponds to post-air puff period with blink artifact. Note (i) reduction in blink artifact magnitude and (ii) reduction in the variability of the pre-air puff pupil trace as training progresses. **(c)** GCA weight estimates. While the highest SNRs elicited clear pupil responses early in training (top), responses to lower SNRs were not clearly graded. Towards the end of the training period, responses to lower SNRs became significant. Bar colors as in Fig. 4. Asterisks correspond to p<0.01 (hypothesis test for linear model coefficients, exact p-values in Supplementary Tables 6 and 7). **(d)** Early in training, pupil responses were not reliable for lower SNRs, resulting in a small percent of trials with significant pupil dilation. Late in training, pupil responses were more reliable, and the percent of pupil responses gradually increased as a function of SNR. Colors are: gray – first 4 days of training, green – next 3 days of training, red – final 3 days of training.

In the early phase, we observed that only deviants with high SNRs elicited a pupil response. Although we averaged over a similar number of deviant trials in the early phase as the experiment shown in Fig. 4, we noticed that pupil responses were significant at few SNRs (Fig. 5c, *top*; Supplementary Table 6), and did not scale with SNR as earlier (Fig. 5d, gray). We only observed significant pupil responses for 20% of trials at most SNRs. We attributed these weak responses to the fact that we ran extended experimental sessions, and deviants occurring later in the session likely evoked little or no pupil responses. In the late training phase, however, we observed that pupil responses were more consistent across SNRs (Fig. 5b, right; note that gray shading, corresponding to ±1 s.e.m., is narrower), a higher proportion of responses were statistically significant (Fig. 5c, bottom; Supplementary Table 7), and both the magnitude of the pupil response as well as the fraction of significant trials scaled gradually with SNR (Fig. 5d, red). Finally, consistent with the earlier thresholds obtained without using an air puff, the detection threshold decreased to around 0 dB SNR. Interestingly, while in the early phase, a large eye blink artifact was visible immediately following the air puff (Fig. 5b, orange shading), in the late phase, this artifact was greatly attenuated, suggesting that the animals adapted to the air puff. Together, these data demonstrate that air puff training can speed up data acquisition and increase the consistency and reliability of pupil responses.

### Pupillometric estimates of call categorization-in-noise thresholds

To probe categorization using an oddball paradigm, we presented calls of one category (*chuts*) embedded in noise at a given SNR as standards (Fig. 6a), and calls of another category (*purrs*) embedded in noise at the same SNR as deviants (Figs. 6b, c). We tested one SNR level per session. But importantly, to ensure that within-category variability was represented in both standard and deviant stimuli, on each standard or deviant trial, calls were randomly chosen from a list of 8 exemplar calls belonging to the appropriate category (Supplementary Fig. 1). Thus, while successive standard stimuli were identical or acoustically indistinguishable in the previous experiments, in the categorization experiment, acoustic changes were present both between successive presentations of standards, as well as between standards and deviants. If low-level acoustic changes were responsible for driving pupil changes, we would expect to see larger responses to the standards themselves, and smaller differences between standard and deviant stimuli. However, if pupil changes also reflected higher-level processes such as categorization, we would expect to see similar trends as in earlier experiments – a small or no response to standards, and large responses to deviants.

**Figure 6:**
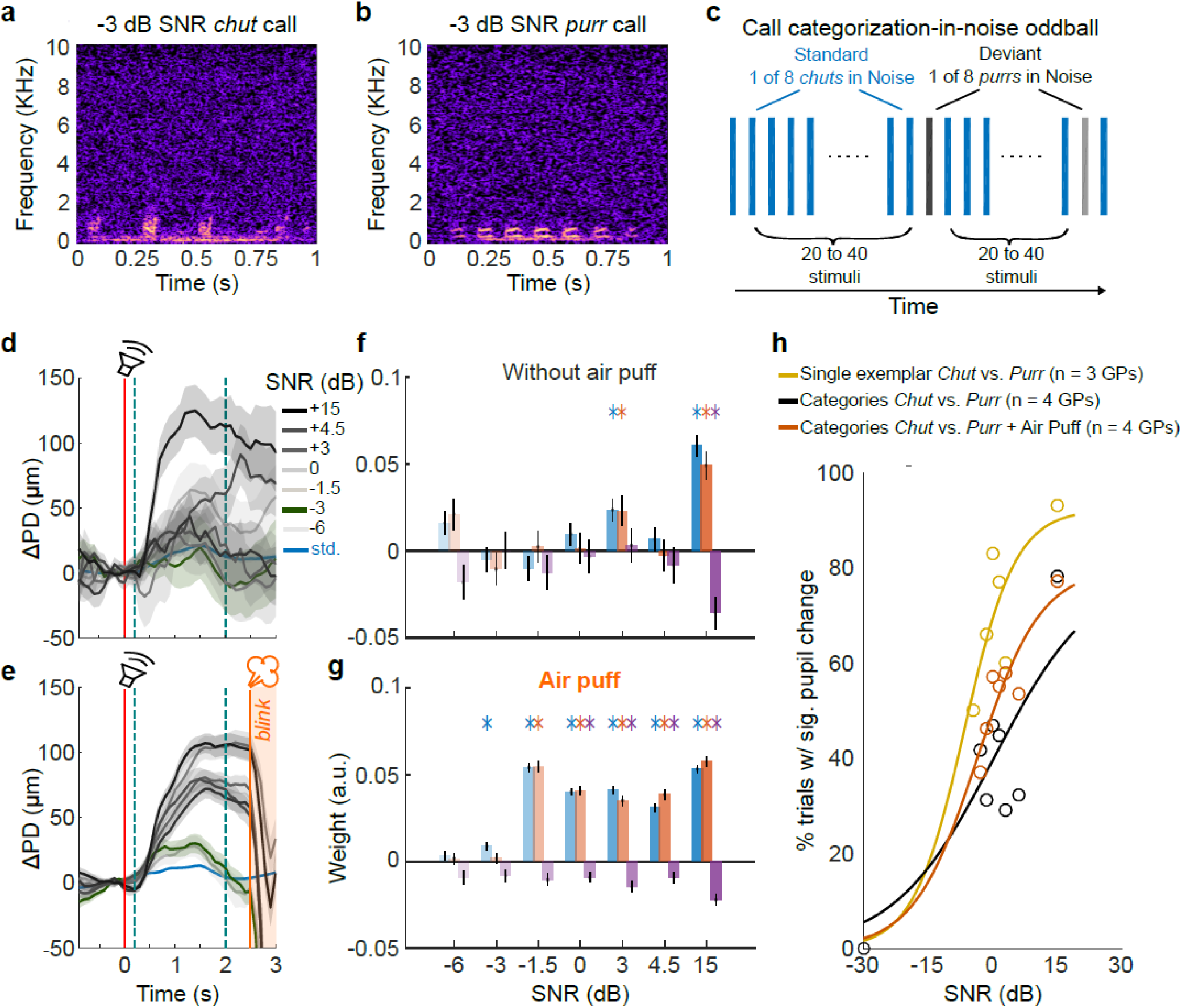
Pupillometric estimates of call categorization-in-noise thresholds. **(a)** and **(b)** Spectrograms of exemplar *chut* and *purr* calls-in-noise at -3 dB SNR. **(c)** For probing call categorization, standard stimuli consisted of *chut* calls randomly drawn from eight exemplars. Deviant stimuli consisted of *purr* calls randomly drawn from eight exemplars. In each experimental session, we added white noise at a different SNR level. **(d)** Average pupil responses from 4 animals. Blue line and shading corresponds to mean and ±1 s.e.m. of pupil responses to standard *chuts*. Gray lines and shading correspond to mean and ±1 s.e.m. of pupil responses to deviant *purrs*. Color gradient corresponds to SNR. Green line and shading denote pupil trace at threshold SNR (−3 dB SNR). Red line corresponds to stimulus onset; teal lines denote GCA window. Without air puff training, we observed a clear pupil response at only the highest SNRs. **(e)** Same as **(d)** but after a week of air puff training (average of training days 8-10 shown). Orange line is air puff onset; orange shading corresponds to post-air puff period with blink artifact. **(f)** GCA weight estimates without air puff training and **(g)** after air puff training. Colors as earlier. Asterisks correspond to p<0.01 (hypothesis test for linear model coefficients, exact p-values in Supplementary Tables 8 and 9). **(h)** Before air puff training (black), the percent of trials with significant responses did not gradually change with SNR. After air puff training (orange), responses were graded with SNR. Yellow line corresponds to a change-detection paradigm where a single exemplar *chut* and *purr* were used as the standard and deviant respectively.

Data from three GPs supported the latter possibility – standard responses were small, and deviant responses were significantly larger than the standard, and of a similar magnitude as those observed in the earlier experiments (Figs. 6d, e). Similar to the call-in-noise experiments, while a large and statistically significant response was observed for ‘clean’ calls without an air puff, neither the magnitudes of other responses nor the fraction of significant responses were correlated with SNR levels (Fig. 6d, f; black points and line in Fig. 6h; Supplementary Table 8). After a week of air puff training, however, we observed robust pupil responses at most SNR levels (Fig. 6e, g; data averaged overs days 8-10 of air puff training shown). Similar to call-in-noise detection, response magnitudes as well as the fraction of trials with significant responses were better correlated with SNR, and responses to SNRs above -3 dB were statistically significant (Fig. 6e, g; orange points and line in Fig. 6h; Supplementary Table 9).

Although the estimated detection (Figs. 4, 5) and categorization (Fig. 6) thresholds were similar, we found that lower thresholds were possible for ‘easier’ versions of the oddball paradigm, where we used a single exemplar of a *chut* and *purr* embedded in white noise as the standard and deviant respectively. In this version, we observed significant responses even at the lowest SNR tested (−6 dB), with the psychometric curve shifted to the left (yellow points and line in Fig. 6h) relative to both the detection and categorization curves. Together, these data indicate that even in the face of standard-to-standard variability, pupillometry can be used to reliably estimate thresholds for tasks that might involve higher-level processes such as categorization.

### Pupillometry as a translatable metric of auditory detection across species

One advantage of pupillometry is that it is an easily-translatable metric that can be acquired from human and animal subjects, across the lifespan and from healthy as well as clinical populations. To demonstrate one example of a complex task for cross-species comparisons, we adapted an auditory figure-ground segregation task that has been previously used in human subjects [36–38]. Standard ‘ground’ stimuli consisted of random 50 ms -long chords with randomly chosen frequency components (Figs. 7a, c). Deviant ‘figure’ stimuli contained about equal acoustic energy as the standards, but with individual frequency components redistributed such that energy was present throughout the stimulus duration within some frequency bands. For example, the ‘figure’ stimulus in Fig. 7b has a coherence value of 6, which means that 6 frequency bands were always ‘on’.

**Figure 7:**
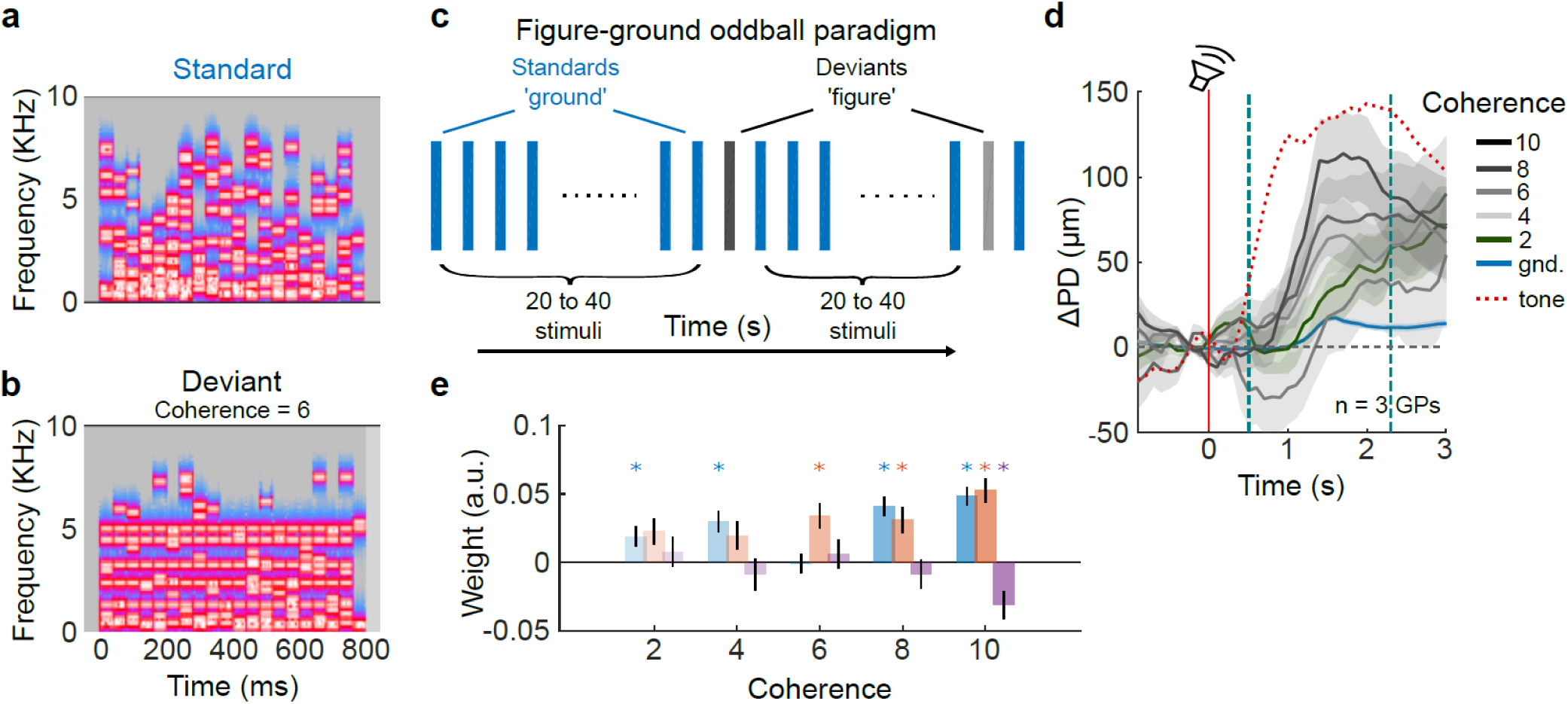
Auditory figure-ground segregation. **(a)** An example standard stimulus consisting only of a background tone cloud (‘ground’ stimulus). **(b)** Deviant stimulus containing an auditory figure embedded in the background, with a coherence of 6. **(c)** The stimulus paradigm consisted of randomly generated standard stimuli containing only of ‘ground’ stimuli, and randomly generated deviants consisting of an auditory figure embedded in the background. One coherence value was probed per experimental session. **(d)** Mean (gray lines) pupil responses evoked by the presence of an auditory figure. Color gradient corresponds to figure coherence. Green line and shading denote pupil trace at threshold (coherence = 2). Shading corresponds to ±1 s.e.m. Blue lines and shading correspond to ground stimuli. Red line is stimulus onset; teal lines correspond to GCA window. Note that the latency of pupil responses in this paradigm was higher compared to other paradigms – red dotted line corresponds to the mean response to the largest harmonic tone deviant from Fig.3. **(e)** GCA weight estimates were significantly different from the standard even for the smallest coherence value tested. Colors as earlier. Asterisks correspond to p<0.01 (hypothesis test for linear model coefficients, exact p-values in Supplementary Table 10).

In three GPs, we found that standard ‘ground’ stimuli evoked little to no pupil responses, whereas deviant ‘figure’ stimuli evoked strong pupil responses that were significantly different from the standard at all coherence values tested (Figs. 7d, e; Supplementary Table 10). The minimum coherence value (coherence = 2) at which we observed a response was consistent with thresholds reported in humans for long stimulus durations [36,37]. Pupil response magnitude was graded as a function of deviant coherence. Interestingly, compared to the earlier detection and categorization paradigms, we noticed that the latency of pupil responses in this paradigm was longer by about 0.5 s (the GCA window was adjusted for this difference). In Fig. 7d, the dotted red line shows the pupil time course for the largest deviant in the harmonic tone discrimination task. This longer latency might reflect two underlying phenomena: 1) the integration time required to detect the change in stimulus statistics between ground and figure stimuli, or 2) the involvement of top-down processes in figure-ground segregation. These data demonstrate the feasibility of using pupillometry to establish equivalencies between human and animal subjects even for sophisticated experimental paradigms.

## Discussion

Our data demonstrate that pupillometry can be used to reliably estimate detection and discrimination thresholds for a wide range of auditory stimuli in untrained animals. Combined with mildly aversive air puff training, threshold estimates can be acquired in about two weeks, with 1 – 2 hours of experimental sessions per animal per day. Because minimal experimenter intervention is required during each session, pupillometry setups are scalable, enabling parallel data acquisition from multiple animals. Although animal habituation constrains the number of deviant trials that can be acquired per day, the oddball paradigm carries advantages when combined with other experimental techniques. For example, the internal state of the animal fluctuates endogenously from trial to trial, and this can be observed in the PD signal as both a change in the baseline PD as well as in the magnitude of the standard-triggered pupil response. This information could be used to sort simultaneously acquired EEG or intracranial electrophysiological data to characterize the state dependence of the neural processing of stimuli [39]. Finally, threshold estimates can be obtained for a variety of stimuli from the same animals without retraining. Because estimated thresholds are consistent with those obtained using operant training, pupillometry could prove highly efficient for large experimental groups. Thus, using pupillometric measures as part of a behavioral testing battery carries numerous advantages.

What does the pupil dilation signal represent? Broadly, in cases where top-down mechanisms are engaged, such as when exerting increased listening effort during active listening in noisy conditions, pupil dilation could be attributed to top-down phenomena that result in increased arousal. In the context of the auditory oddball paradigm, it is likely that the pupil dilation signal reflects bottom-up capture of attention by the deviant stimulus. It is thus important to understand whether PD changes are merely triggered by any acoustic change, or if the PD signal can also be modulated by prior experience, top-down signals, and the effects of training or learning. Our data suggests that not all acoustic changes are equal – negative changes (such as the occurrence of a gap in noise) fail to evoke pupil dilation, and pupil dilation increases proportionally with acoustic distance between the standard and deviant stimuli. Pupil dilation may also not be purely driven by low-level acoustic changes – the pupil response is tolerant to within-category variations, and shows robust responses to changes in categorical identity. Finally, pupil responses can be shaped through training. These data suggest that the estimates of perceptual boundaries that are obtained using pupillometry reflect an integrative effect of stimulus differences and internal state of the animal. But the question remains open as to whether the relative influence of endogenous versus exogenous factors on pupil diameter differs by species. Being a prey species facing predation pressure, arousal in GPs may be more inclined to be affected by small external sensory changes, leading to the large and scaled PD changes that we have observed.

How do our pupillometric threshold estimates compare to estimates obtained using operant training? First, we estimated F0 discrimination threshold at 0.75 semitones (at F0 = 440 Hz), or a 4.5% increase in frequency. Here, we conservatively estimated the threshold as the ΔF (or SNR for other tasks) required to reach the half-maximum point of the psychometric fits because this value also corresponded to the point at which GCA coefficients were statistically significant. Few studies have measured frequency discriminability in GPs, primarily because training GPs in operant tasks is time-consuming. Using aversive conditioning in a Go/No Go stimulus paradigm, Heffner et al. [40] had estimated the pure tone difference thresholds in two GP subjects to be a 3.5% change. More recently, in marmosets (go/no go task), frequency difference thresholds for pure tone discrimination was estimated to range from about 3 semitones at low frequencies, to 0.37 semitones at marmoset call frequencies [41]. Our pupillometric threshold estimate of 0.75 semitones (4.5%) for harmonic complexes is in agreement with these values. For resolved harmonic complexes with F0 of 440 Hz, the difference threshold at F0=440 Hz was estimated at 0.37 semitones in marmosets [42], about half our estimate of 0.75 semitones in GPs. Keeping in mind the species differences, our pupillometric estimates thus seem consistent with those obtained using conventional operant training techniques. Second, we estimated call- and tone-in-noise detection thresholds at about -1.5 dB SNR. Our threshold estimate lies well within the range of values reported for other stimulus-in-noise detection tasks (range: -2 dB SNR to +5 dB SNR) obtained using operant conditioning in a variety of species [43–47]. While aversive conditioning (such as an electric shock) seems to result in lower thresholds, our pupillometric estimates are in closer agreement with threshold values obtained using appetitive conditioning (water or food reward). These results are also consistent with earlier observations in barn owls, where pupillometric estimates closely matched those obtained using operant conditioning [23]. Recent observations in human subjects also confirmed that pupillometric estimates of tone detection thresholds were in close agreement with estimates based on subject reports [29]. While operant conditioning remains the gold standard for evaluating perceptual thresholds, our results suggest that pupillometry can be used to obtain close estimates of these thresholds efficiently.

In conclusion, we have demonstrated that pupillometry can be used to reliably estimate thresholds for various auditory behaviors in animal models. Because pupillometric data can be acquired reasonably quickly in untrained animals using multiple stimulus paradigms, this technique could be especially useful for obtaining high-throughput behavioral threshold estimates from large experimental groups. Finally, the non-invasive nature of pupillometry makes acquiring comparative data between human and animal subjects relatively easy. Combined with other non-invasive comparative measurements such as EEG recordings, this sets the stage for using harmonized stimulus and data acquisition protocols to establish broad behavioral similarities between human and animal subjects, which can then be followed up in animals using invasive recordings to interrogate the neural bases of these behaviors. Because pupillometric signals are affected in some clinical populations [22,30–32], our study sets the stage for using pupillometry as an additional phenotype for potential animal models of these disorders.

## Methods

### Animals

All experimental procedures conformed to NIH Guide for the care and use of laboratory animals, and were approved by the Institutional Animal Care and Use Committee (IACUC) of the University of Pittsburgh. We acquired data from 5 male and 6 female adult, wild-type, pigmented guinea pigs (Elm Hill Labs, Chelmsford, MA), weighing ∼600-1000 g over the course of the experiments. Some animals also participated in unrelated electrophysiological studies.

### Surgical procedures

All experiments were conducted in unanesthetized, head-fixed, passively-listening animals. To achieve head fixation, a custom head post and recording chambers were first surgically anchored onto the skull using dental acrylic (C&B Metabond, Parkell Inc., Brentwood, NY) following aseptic techniques under isoflurane anesthesia. Post-surgical care, including systemic and topical analgesics, was provided for 3 – 5 days. Following a 2-week recovery period, animals were gradually adapted to the experimental setup by slowly increasing the duration of head fixation.

### Pupillometry data acquisition

All experiments were performed in a walk-in double-walled sound attenuating chamber (IAC), with the inner walls covered with anechoic foam (Sonex One, Pinta Acoustic, Minneapolis, MN). Animals were placed in a custom acrylic enclosure fixed to a vibration-isolation tabletop that provided loose restraint of the body. The enclosure allowed the animal to make small movements such as postural adjustments during experiments, but not large movements. A piezoelectric sensor (SEN-10293, SparkFun Electronics, Niwot, CO) placed beneath the enclosure was used to detect and record these movements. The head post was affixed to a rigid frame, providing stable head-fixation with the animal facing a loudspeaker (W4-1879 full-range driver, Tang Band Speaker, Taipei, Taiwan) mounted on the wall of the sound attenuating chamber at a 90 cm distance. The frequency response of the loudspeaker was periodically calibrated using a free-field microphone (Type 4940, Bruel & Kjaer, Denmark) and verified to have a flat frequency response (± 5 dB) over the range of guinea pig audible frequencies (0.5 – 32 Khz). Lighting conditions within the sound chamber were kept constant across experimental sessions.

One eye was illuminated with an infrared LED array (MCU902, Arrington Research, Scottsdale, AZ), and video of the pupil acquired using a camera with an infrared filter (MCU902, Arrington Research, Scottsdale, AZ) placed ∼15 cm from the imaged eye. The diameter of the pupil was obtained using a thresholding algorithm (ViewPoint, Arrington Research, Scottsdale, AZ). The imaged eye was illuminated with bright and diffuse white LED lighting, with the brightness adjusted to bring the baseline PD to ∼3.5 mm. This ensured that baseline lighting conditions were consistent both across sessions and across animals, and allowed for sufficient dynamic range to observe sound-evoked pupil dilation. Pupil diameter and pupil velocity were acquired at a sampling rate of 90 Hz and converted to an analog voltage output (PCI-DDA02/12, Measurement Computing Corporation, Norton, MA). All data were recorded at 1 KHz using a neural interface processor’s (Scout, Ripple Neuro, Salt Lake City, UT) analog and digital inputs to synchronize stimulus delivery times, the PD trace, the motion trace from the piezo sensor, and air puff delivery times.

### Auditory Stimuli

All stimuli were generated in Matlab (ver. 2018a, Mathworks, Inc., Natick, MA) at a sampling rate of 100 KHz, converted to analog (PCI-6229, National Instruments, Austin, TX), attenuated (PA5, Tucker-Davis Technologies, Alachua, FL), power-amplified (SA1, Tucker-Davis Technologies, Alachua, FL), and delivered through a calibrated speaker (W4-1879 full-range driver, Tang Band Speaker, Taipei, Taiwan). For most experiments, we used an auditory oddball paradigm with 0.5 s or 1 s long ‘standard’ stimuli that occurred with high probability (>90%). ‘Deviant’ stimuli of equal length were interspersed rarely among the standard stimuli. Stimuli occurred with high temporal regularity (every 3 seconds), with inter-stimulus interval chosen to allow pupil responses to reach their maxima. To minimize possible effects of decreasing pupil dilation magnitudes over the course of a session, we employed a Latin square design to counterbalance deviant order across experimental sessions. Each of the deviant stimuli occurred at a unique position in the stimulus sequence order in each experimental session. Thus, across sessions, possible changes in pupil magnitude as a result of within-session adaptation would be averaged out. To obtain a complete data set from each animal, we used eight different deviant stimuli presented in unique orders over eight sessions. To avoid adaptation, we limited experimental sessions to about 30 min. per day, during which data for one stimulus order from two stimulus paradigms could be acquired. We assessed how many deviant trials were obtained in a given session without interference from blinking and motion-related pupil dilation. Some sessions were repeated if more than half the number of deviant trials were discarded. All stimuli were presented at 70 dB SPL. The identity of the standard and deviant stimuli varied with the experimental paradigm being tested, described below.

### Harmonic complexes

The standard stimulus was a 1-second-long harmonic complex with fundamental frequency (F0) of 440 Hz and consisting of the first sixteen harmonic components. Deviant stimuli were 1s long and also contained 16 harmonic components, with parametrically varied F0s (0.25 to 2 semitones greater than the standard F0).

### Call-in-noise

We used ‘*purr*’ and ‘*chut*’ guinea pig vocalizations that we recorded by placing animals inside a sound-attenuated enclosure. All vocalizations were recorded in our animal colony using Sound Analysis Pro [48] by placing one or more animals in a sound-attenuated booth and by recording vocalizations using a directional condenser microphone (C-2, Behringer). Two observers manually segmented and classified vocalizations into categories based on previously published descriptions [49,50]. The calls were normalized by their r.m.s. amplitudes. Depending on recording quality, the signal-to-noise ratios (SNRs) of these recorded calls were estimated at 15 – 30 dB SNR. Standard stimuli consisted of 1 s long white noise samples, generated independently for each standard stimulus. Deviant stimuli consisted of calls embedded in white noise at different SNR levels. To generate call-in-noise stimuli at different SNRs, we added white noise of equal length to the calls (gated noise) such that the signal-to-noise ratio, computed using r.m.s. amplitudes, varied between -4.5 dB and +6 dB SNR. This range of SNRs was chosen to maximize sampling over the linear part of psychometric curve fits. As a control stimulus, we also generated harmonic complex in noise stimuli. All stimuli were presented at an r.m.s. amplitude of 70 dB SPL.

### Call category discrimination

Eight different exemplars of *chut* and *purr* vocalizations were r.m.s.-equalized, and white noise at different intensities was added as described earlier to generate call-in-noise stimuli. We acquired data corresponding to one SNR level during each experimental session. In each session, eight different *chut* vocalizations at a particular SNR were used as standards, and eight different *purrs* at the same SNR were used as deviant stimuli. To compare categorization results against an easier version of the task, we also gathered pupil data in response to an oddball paradigm with a single example each of the *chut* and *purr*.

### Figure-ground segregation

Stimuli were generated based on previous studies [36–38]. Briefly, standard ‘ground’ stimuli and deviant stimuli containing a ‘figure’ were 1 s long, and consisted of random chords made of 50 ms tone pips. Stimuli spanned a five-octave frequency range (200 – 6400 Hz). Stimulus density was set at 1 tone pip per octave per chord. To generate deviant ‘figure’ stimuli at different coherences, we randomly chose n frequency bins (where n is the coherence value), and added tone pips at those frequencies in all time bins. We added an equal number of tone pips to all time bins of the standard ‘ground’ stimuli, but at random frequencies, to maintain constant tone pip density across stimuli. We tested five coherence values in our experiments (one per session).

### Air puff training

In the experiments described in Figs. 2e, 5, and 6, we delivered a brief (100 ms) air puff 2.5 s after the onset of the deviant stimulus. This delay in air puff delivery was chosen to allow the PD to reach a peak dilation uninterrupted by the air puff. Air puffs were delivered from a pipette tip placed ∼15 cm in front of the animal directed at the animal’s snout. The pipette tip was connected using silicone tubing via a pinch valve (EW98302-02, Cole-Palmer Instrument Co., Vernon Hills, IL) to a regulated air cylinder, with the air pressure regulated to be between 20 and 25 psi. The timing and duration of the air puffs were controlled by activating the pinch valve using custom computer-controlled electronics. To track training, we performed these experiments using identical stimulus conditions for ten days.

### Analysis and Statistics

All analysis was performed using custom code written in Matlab (ver. 2018a, Mathworks Inc., Natick, MA). Specific functions and toolboxes used are mentioned where applicable below.

### Motion detection and trial exclusion

First, we extracted and detrended the motion trace over the entire session (∼12 minutes) and measured the standard deviation (SD) of the motion trace. We then obtained the times of motion trace peaks that crossed a threshold of 5 SDs and were separated from other peaks by at least one second (Fig. 1b, bottom). Because the motion-induced pupil trace peaked ∼2 s after a motion event and slowly returned to baseline (Fig. 1c), we discarded any trials that occurred within 7 seconds of a motion event to allow motion-related changes to subside and for stimulus-evoked changes to be detected reliably.

### Data pre-processing

We first linearly detrended the PD trace and converted the units from voltage to micrometers. We detected eye blinks, defined as PD changes exceeding 400 μm/ms, and removed them by linearly interpolating the PD trace in a 200 ms time window centered at the detected blink time (note that GPs do not blink frequently). We then downsampled the data to 10 Hz. We computed the average baseline PD for each stimulus in a 500 ms window just prior to stimulus onset. We then extracted PD traces in a window beginning 1 s before stimulus onset and lasting 3 s after stimulus offset, and subtracted the baseline PD from these traces. Pupil traces for each stimulus condition were then averaged across sessions and animals.

### Quantification of stimulus-related PD changes

To quantify the fraction of pupil dilation events that could be explained by stimulus presentation and animal motion, we smoothed the pupil trace over each experimental session (50 ms rectangular window) and detected peaks (‘findpeaks’ function in Matlab) with: 1) a prominence of at least 25 μm, 2) inter-peak separation of at least 500 ms, and 3) a halfwidth of at least 500 ms. We then calculated the fraction of these peaks which occurred within 2 s of a stimulus presentation or a motion event. The average number of pupil dilation events, and the fraction associated with stimuli and motion were computed over seven GPs, using data from the harmonic tone discrimination paradigm (Fig. 3).

### Analysis

We adopted two analytical approaches to quantify PD changes. First, we built distributions of peak PDs in a 1-second-long analysis window beginning 1 s after stimulus onset. For each deviant trial, we determined if the distribution of peak PD values was significantly different from the pooled distribution of PD values obtained for all standard stimuli using single-tailed two-sample t-tests. We thereby quantified, for each deviant, the fraction of trials that showed statistically significant increases in PD compared to standard stimulus trials. To these data, we used the ‘fitnlm’ Matlab function (Statistics toolbox) to fit psychometric functions of the form [51]:

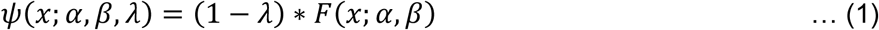

where F is the Weibull function, defined as 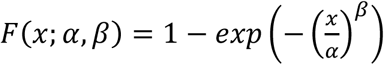, α is the shift parameter, β is the slope parameter, and λ is the lapse rate. This analysis quantified, for each deviant, how *reliably* PD changes could be elicited from trial to trial.

### Growth curve analysis (GCA)

To complement the above analysis, we used GCA [13,35] to quantify the *magnitude* and *shape* of stimulus-evoked PD changes. While typical analyses of pupil dilation (as above) focus on PD changes evoked in experimenter-defined analysis windows, this approach could potentially lead to: 1) experimenter bias in picking the analysis window, 2) multiple-comparisons concerns, and 3) a loss of statistical power because the averaging of PD over the entire window disregards the temporal evolution of the pupil trace. To overcome these issues, we fit linear mixed-effect models with subject-level intercepts as random effects, and orthogonal time polynomials of up to order two as fixed effects, with each deviant treated as a separate group, to the rising phase of the pupil trace (200ms post stimulus onset to 2 seconds post stimulus onset in Figs. 3 – 6; 500 ms post onset to 2.3 s post onset in Fig. 7). The rising phase of the pupil trace was modeled using the following formula [13]:

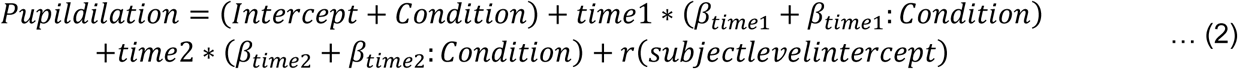

where *time1* and *time2* correspond to orthogonal linear and quadratic time polynomials, and βs correspond to weights. Estimates of mean weights (βs) and their standard errors were obtained using the ‘fitlme’ function in MATLAB. Statistical significance was evaluated using linear hypothesis tests for linear regression model coefficients (‘coefTest’ in the Matlab Statistics Toolbox), employing Satterthwaite’s correction for large degrees of freedom. We restricted fixed effects to time polynomials of order two because the addition of a third-order polynomial did not result in a significant increase in model performance, and third-order weights were not statistically significant. In this formulation, the β value for the intercept roughly corresponds to the mean pupil dilation in the analysis window, the β value for the linear time polynomial (*time1*) corresponds to the slope of the pupil dilation trace, and the β value for the quadratic time polynomial (*time2*) corresponds to the acceleration of the pupil dilation trace. Because of the tendency of the PD to revert to baseline a few seconds after stimulus onset, the quadratic weights were often observed to be negative.

## Data Availability

The datasets generated and/or analyzed during the current study will be made available upon reasonable request to the corresponding author.

## Acknowledgements

We thank Madelyn McAndrew and Dr. Marianny Pernia for assistance with performing pupillometry experiments and animal care, and Samuel Li for assistance with recording and categorizing vocalizations. We thank Dr. Yi Zhou (Arizona State University) for insightful comments on the manuscript. We thank Jacie McHaney and Dr. Bharath Chandrasekaran for suggesting growth curve analysis to evaluate pupil dilation data. We thank Stacy Cashman and Mark Petts for surgical support; Dr. Amanda Fisher for veterinary support; and Jillian Harr, Sarah Gray, Julia Skrinjar, Brent Barbe, and Elizabeth Chasky for animal care. SS is grateful for support from the NIH (R01DC017141), a Pennsylvania Lions Hearing Research Foundation grant, and a NARSAD Young Investigator award from the Brain and Behavior Research Foundation.

## Author Contributions

S.S. and P.M-L. designed the experiments. P.M-L., M.K., and I.K. performed all pupillometry experiments. S.S. analyzed the data with inputs from M.K. and P.M-L. P.M-L. and S.S. prepared manuscript figures, and S.S. wrote the manuscript with inputs from all co-authors.

