## Supplemental Figure and Tables 1-10 for "Pupillometry as a reliable metric of auditory detection and discrimination across diverse stimulus paradigms in animal models"

**Supplementary Information contains:**

1 Supplementary Video (MP4)

1 PDF file with:

- Legend for Supplementary Video
- Supplementary Fig. 1 and Legend
- Supplementary Discussion
- 10 Supplementary Tables containing exact p-values for all statistical tests used in the main manuscript.

**Legend for Supplementary Video 1: Pupil diameter changes in an auditory oddball task.**
Top panel: Video of GP eye under infrared illumination (2x speed).
Middle panel: White line corresponds to the pupil diameter trace. X-axis numbers are time in seconds. Gray bars correspond to white noise standards. Red bar corresponds to the deviant stimulus (in this case, a clean GP call).
Bottom panel: Voltage output of the piezoelectric sensor. Slower deflections correspond to respiration, rapid deflections in the later part of the trace corresponds to postural shifts made by the animal. Yellow line is time marker.

**Supplementary Figure 1: Spectrograms of guinea pig vocalizations used in pupillometry experiments. (A)** Chut vocalizations (top) and Purr vocalizations (bottom) used in the experiments described in Figs. 4 – 6. **(B)** Chuts (top) and Purrs (bottom) with white noise added to result in a final signal-to-noise ratio of -3 dB SNR.

(Figure on next page)

**Supplementary Fig. 1**

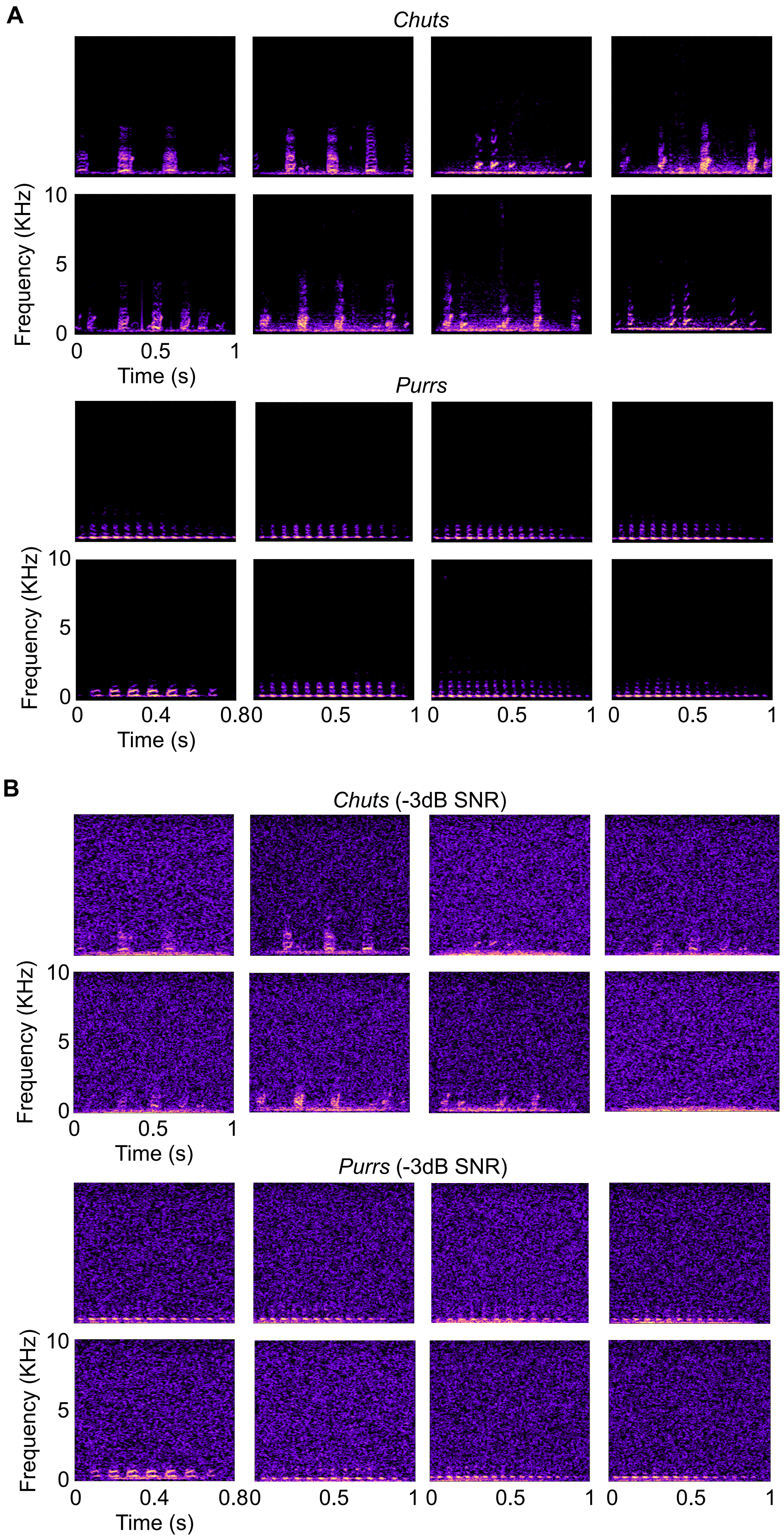

**Supplementary Discussion**

A minor methodological disadvantage is that in our experiments, animals were first surgically implanted with a headpost for head-fixation. Requiring head-posted animals potentially limits the scalability of using pupillometry for high-throughput evaluation of large experimental groups. While pupil tracking is possible in freely-moving animals, these systems also require some form of head-anchoring [1,2], and because pupil dilation accompanies animal motion [3,4], a free-moving preparation is a sub-optimal solution. Headpost-free restraints are possible – for example, for functional magnetic resonance imaging in small animals, 3d-printed ‘helmets’ have been used for headpost-free head fixation [5]. We believe that pupillometry with non-invasive restraint in non-headposted animals, using recent advances in markerless pose-estimation methods [6,7] to accurately track PD despite some head motion, is an easily achievable solution for acquiring high-throughput and high-quality pupillometry data.


**Supplementary Table 1:** Exact p-values for statistical analysis in Fig. 3E.

Tests: ANOVA (F = 107.98, dF = 8, p = $1.63 \times 10^{-180}$), followed by Bonferroni-corrected post-hoc tests.

| **(Baseline vs. ) ΔF** | **p-val.** |
|---|---|
| 0.25 | 0.072 |
| 0.50 | 1 |
| 0.75 | $6.2 \times 10^{-7}$ |
| 1.00 | $3.1 \times 10^{-11}$ |
| 1.25 | $1.4 \times 10^{-31}$ |
| 1.50 | $4.1 \times 10^{-11}$ |
| 1.75 | $2.8 \times 10^{-74}$ |
| 2.00 | $3.2 \times 10^{-54}$ |

**Supplementary Table 2:** Exact p-values for statistical analysis in Fig. 3H.

<u>Tests:</u> Linear hypothesis tests for model coefficients with Satterthwaite's correction for large degrees of freedom. For brevity, only significant p-values < 0.01, marked with asterisks in Fig. 3H, are shown.

| ΔF (semitones) | Intercept p-val. | Time1 p-val. | Time2 p-val. |
|---|---|---|---|
| 0.75 | $9.2 \times 10^{-8}$ | 0.0025 | |
| 1.00 | $3.5 \times 10^{-8}$ | 0.0057 | |
| 1.25 | $4.6 \times 10\text{-}^{24}$ | $4.8 \times 10^{-8}$ | |
| 1.50 | $3.7 \times 10^{-11}$ | | |
| 1.75 | $6.0 \times 10^{-43}$ | $1.0 \times 10^{-23}$ | $5.1 \times 10^{-5}$ |
| 2.00 | $1.0 \times 10^{-35}$ | $3.6 \times 10^{-16}$ | $6.7 \times 10^{-4}$ |

**Supplementary Table 3:** Exact p-values for statistical analysis in Fig. 4F (*top*) *Chut* calls-in-noise.

<u>Tests:</u> Linear hypothesis tests for model coefficients with Satterthwaite's correction for large degrees of freedom. For brevity, only significant p-values < 0.01, marked with asterisks, are shown.

| SNR (dB) | Intercept p-val. | Time1 p-val. |
|---|---|---|
| -4.5 | $1.6 \times 10^{-7}$ | 0.0019 |
| -1.5 | $1.3 \times 10^{-5}$ | $1.1 \times 10^{-4}$ |
| 1.5 | $6.0 \times 10^{-14}$ | $7.9 \times 10^{-12}$ |
| 3.0 | $6.7 \times 10^{-13}$ | $2.0 \times 10^{-6}$ |
| 6.0 | $7.4 \times 10^{-34}$ | $1.5 \times 10^{-16}$ |
| 20.0 | $1.2 \times 10^{-38}$ | $3.6 \times 10^{-25}$ |

**Supplementary Table 4:** Exact p-values for statistical analysis in Fig. 4F (*mid*) *Purr* calls-in-noise.

<u>Tests:</u> Linear hypothesis tests for model coefficients with Satterthwaite's correction for large degrees of freedom. For brevity, only significant p-values < 0.01, marked with asterisks in Fig. 4F, are shown.

| SNR (dB) | Intercept p-val. | Time1 p-val. | Time2 p-val. |
|---|---|---|---|
| -3.0 | $3.8 \times 10^{-9}$ | 0.0017 | |
| 0.0 | $2.6 \times 10^{-12}$ | $1.8 \times 10^{-11}$ | |
| 1.5 | $1.2 \times 10^{-10}$ | $2.3 \times 10^{-9}$ | |
| 3.0 | 0.0016 | | |
| 20.0 | $3.3 \times 10^{-80}$ | $5.3 \times 10^{-46}$ | $4.8 \times 10^{-6}$ |

**Supplementary Table 5:** Exact p-values for statistical analysis in Fig. 4F (*bottom*) harm. tone-in-noise.

<u>Tests:</u> Linear hypothesis tests for model coefficients with Satterthwaite's correction for large degrees of freedom. For brevity, only significant p-values < 0.01, marked with asterisks in Fig. 4F, are shown.

| SNR (dB) | Intercept p-val. | Time1 p-val. |
|---|---|---|
| -4.5 | $3.0 \times 10^{-4}$ | 0.0053 |
| -3.0 | $1.0 \times 10^{-5}$ | |
| 0.0 | $2.4 \times 10^{-4}$ | 0.0024 |
| 1.5 | 0.0034 | |
| 3.0 | $1.8 \times 10^{-26}$ | $2.4 \times 10^{-6}$ |
| 6.0 | $6.3 \times 10^{-12}$ | $3.2 \times 10^{-9}$ |
| 20.0 | $4.0 \times 10^{-7}$ | $1.1 \times 10^{-4}$ |

**Supplementary Table 6:** Exact p-values for statistical analysis in Fig. 5C (*top*), call-in-noise detection for Days 1-4 of air puff conditioning.

<u>Tests:</u> Linear hypothesis tests for model coefficients with Satterthwaite's correction for large degrees of freedom. For brevity, only significant p-values < 0.01, marked with asterisks in Fig. 5C, are shown.

| SNR (dB) | Intercept p-val. | Time1 p-val. | Time2 p-val. |
|---|---|---|---|
| 1.5 | $2.5 \times 10^{-4}$ | | 0.0039 |
| 3 | $1.5 \times 10^{-14}$ | $1.9 \times 10^{-4}$ | |
| 20 | $1.2 \times 10^{-22}$ | $6.8 \times 10^{-15}$ | 0.0011 |

**Supplementary Table 7:** Exact p-values for statistical analysis in Fig. 5C (*bottom*), call-in-noise detection for Days 8-10 of air puff conditioning.

<u>Tests:</u> Linear hypothesis tests for model coefficients with Satterthwaite's correction for large degrees of freedom. For brevity, only significant p-values < 0.01, marked with asterisks in Fig. 5C, are shown.

| SNR (dB) | Intercept p-val. | Time1 p-val. | Time2 p-val. |
|---|---|---|---|
| -3 | $2.1 \times 10^{-6}$ | $8.7 \times 10^{-5}$ | |
| -1.5 | 0.0022 | | |
| 1.5 | $2.4 \times 10^{-6}$ | | |
| 6 | $7.9 \times 10^{-9}$ | 0.0022 | |
| 20 | $3.9 \times 10^{-43}$ | $9.9 \times 10^{-12}$ | 0.0083 |

**Supplementary Table 8:** Exact p-values for statistical analysis in Fig. 6F, call categorization-in-noise without air puff conditioning.

Tests: Linear hypothesis tests for model coefficients with Satterthwaite's correction for large degrees of freedom. For brevity, only significant p-values < 0.01, marked with asterisks in Fig. 6F, are shown.

| SNR (dB) | Intercept p-val. | Time1 p-val. | Time2 p-val. |
|---|---|---|---|
| 3 | $2.3 \times 10^{-4}$ | 0.0064 | |
| 15 | $1.7 \times 10^{-23}$ | $7.3 \times 10^{-10}$ | $9.1 \times 10^{-5}$ |

**Supplementary Table 9:** Exact p-values for statistical analysis in Fig. 6G, call categorization-in-noise with air puff conditioning.

Tests: Linear hypothesis tests for model coefficients with Satterthwaite's correction for large degrees of freedom. For brevity, only significant p-values < 0.01, marked with asterisks in Fig. 6G, are shown.

| SNR (dB) | Intercept p-val. | Time1 p-val. | Time2 p-val. |
|---|---|---|---|
| -3 | $1.3 \times 10^{-4}$ | | |
| -1.5 | $1.0 \times 10^{-119}$ | $1.1 \times 10^{-71}$ | 0.0024 |
| 0 | $1.7 \times 10^{-97}$ | $1.4 \times 10^{-58}$ | 0.0012 |
| 3 | $5.4 \times 10^{-83}$ | $1.4 \times 10^{-35}$ | $5.0 \times 10^{-6}$ |
| 4.5 | $7.4 \times 10^{-47}$ | $8.4 \times 10^{-42}$ | 0.0039 |
| 15 | $3.4 \times 10^{-140}$ | $5.7 \times 10^{-96}$ | $3.1 \times 10^{-12}$ |

**Supplementary Table 10:** Exact p-values for statistical analysis in Fig. 7E, figure-ground segregation.

Tests: Linear hypothesis tests for model coefficients with Satterthwaite's correction for large degrees of freedom. For brevity, only significant p-values < 0.01, marked with asterisks in Fig. 7E, are shown.

| Coherence | Intercept p-val. | Time1 p-val. | Time2 p-val. |
|---|---|---|---|
| 2 | $7.9 \times 10^{-4}$ | | |
| 4 | $7.7 \times 10^{-5}$ | | |
| 6 | | $2.2 \times 10^{-4}$ | |
| 8 | $5.6 \times 10^{-9}$ | $7.9 \times 10^{-4}$ | |
| 10 | $3.3 \times 10^{-13}$ | $1.9 \times 10^{-9}$ | 0.0018 |
